# Substance matters: IL5 and IL33 activation of eosinophils on periostin and fibrinogen induce cytoskeletal reorganization and cell death

**DOI:** 10.64898/2026.02.28.708742

**Authors:** Joshua M. Mitchell, Deane F. Mosher

**Affiliations:** Department of Biomolecular Chemistry, University of Wisconsin-Madison, WI; Department of Medicine, University of Wisconsin-Madison; Morgridge Institute for Research, Madison, WI

## Abstract

Human eosinophils activated in suspension with IL5 or IL33 undergo morphological change prior to adhesion. Refractive granules, which contain major basic protein-1 and other toxic proteins, move to one side of the cell, the granulomere, and the two nuclear lobes move to the other. How these features persist when eosinophils become adherent and migrate is not known. We now compare behavior of activated eosinophils on surfaces coated with ITGAM/ITGB2-integrin ligands fibrinogen or periostin using live cell imaging of reporters of tubulin/actin organization and cell viability. We find that unlike eosinophils activated with IL5, IL33-activated eosinophils undergo two stages of activation; a preliminary pear-like activation in which the cell develops polarity, followed by a flattening of the eosinophil into a thin pancake-like morphology with less discrete polarization. IL5-treated eosinophils migrated persistently for more than an hour with nucleopod in the back. In contrast, IL33-treated eosinophils moved more slowly and within 30 min transitioned to a flattened morphology with nuclear lobes in the center and dispersed motile granules. Loss of cell viability after an hour, although variable, in all comparisons was greater among IL33-treated eosinophils on periostin. We sought to understand how cytoskeletal elements may drive these differences in morphology. Cytoskeletal elements had similar responses when activated with IL5/IL33; vimentin collapsed from a web-like network at the periphery of the cell and condensed adjacent to the nucleopod/nuclear interface, f-actin was found in the granulomere as well as the tip of the nucleopod and forward periphery, and microtubules radiated from the microtubule organizing center (MTOC) spanning both the nucleopod and the granulomere. The dynamic formation of microtubules correlated with cellular locomotion, suggesting mesenchymal migration within these cells. These in vitro findings suggest that adhesion plays an important role in determining functional morphology and demonstrates new insights into IL33-activated eosinophils. This work suggests roles for activators and adhesive substrates in regulating the behavior of activated eosinophils in tissues.

## Introduction

Eosinophils are white blood cells important for normal homeostasis and the inflammation that characterizes eosinophilic disorders^1^. Eosinophils contain a large specific granule that contains high concentrations of major basic proteins 1 (MBP1, a fragment of PRG2) and 2 (MBP2, a fragment of PRG3), secretory RNases 2 (RNASE2, also known as eosinophil-derived neurotoxin or EDN) and 3 (RNASE3, also known as eosinophil cationic protein or ECP), and eosinophil peroxidase (EPX) [PMID: **8476270, 2734298, 6644025, 2473157, 11698487, 6281334**]. An important determinant of function is the activation that results in migration from blood into tissues, where eosinophils participate in mediator crosstalk with other cells and may undergo oxidative burst and release granular contents through piecemeal degranulation or cell lysis and extracellular deposition of granules^2–8^.

When studied in suspension, transition from naïve to activated eosinophils is characterized by changes in cell morphology that depend on the activator. IL5-family cytokines cause dramatic changes in cell morphology and polarization with the nuclear lobes moving to one end, the nucleopod, and the granules into the other, the granulomere^9^. Polarization of suspended eosinophils caused by IL33 produce the same phenomena, but is less dramatic^10^.

IL5 is the eosinophil’s “private” cytokine that upregulates eosinophil production and causes eosinophil activation that can lead to deleterious effects on human health^11–13^. Effects of IL33 on eosinophils are not as well characterized as those of IL5, although the significance of IL33 for eosinophil development and related diseases and disorders is becoming increasingly appreciated^11,14–19^.

We previously reported adhesion and random motility and described video phase microscopy of IL5-treated eosinophils placed of surfaces coated with periostin^20–22^, an extracellular matrix protein and ITGAM/ITGB2-integrin ligand that is deposited in tissues undergoing eosinophilic inflammation^23^. In addition, IL33-treated eosinophils have been reported to adhere to periostin^24^. The majority of IL5-treated eosinophils kept the polarized shape adopted in suspension while migrating persistently with the nucleus at the rear, granules continuously being gathered together into the granulomere that was in front of the nucleus, and a ruffling forward edge exploring the substrate surface^22^. On denser coatings of periostin, some eosinophils transiently adopted a flattened pancake morphology with dispersed granules and nuclear lobes. Incubations of IL5-activated eosinophils for several hours on coatings of fibrinogen, also a ITGAM/ITGB2-integrin ligand, has been reported to cause cell lysis^25^.

How the polarized morphology of eosinophils activated in suspension evolves when eosinophils move into tissues needs to be understood in greater depth. We now compare the behavior of eosinophils activated with IL5 or IL33 on substrates of periostin or fibrinogen using fluorescent microscopy of fixed cells and live cell imaging by phase microscopy and fluorescent microscopy of reporters of tubulin and actin organization and cell viability. We note the polarization and dramatic reorganization of activated adherent eosinophils, and the distinct morphologies that develop when activated with IL5 or IL33. We observe that microtubules imaged in live eosinophils as compared to fixed are far more extensive than previously imaged, and that other cytoskeletal components provide crucial support to the nucleus and to the granules for cellular integrity. We also observe unique procession of activation of IL5 compared to IL33, and finally note that loss of nuclear integrity signals imminent cell death of activated eosinophils.

## Results

### Morphology of eosinophils on fibrinogen evolves differently after activation by IL5 or IL33

By confocal phase microscopy, adherent IL5-activated eosinophils maintained an acorn-like morphology and migrated across fibrinogen similarly to what we described for IL5-activated eosinophils on periostin^9,22^ (**Figure 1**). Most cells remained migratory for up to 90 min. IL33-activated eosinophils behaved differently. Initially adherent cells assumed a polarized state that was blunted and lacked the striking nucleopod tip of IL5-activated eosinophils (**Figure 1**).

**Figure 1:**
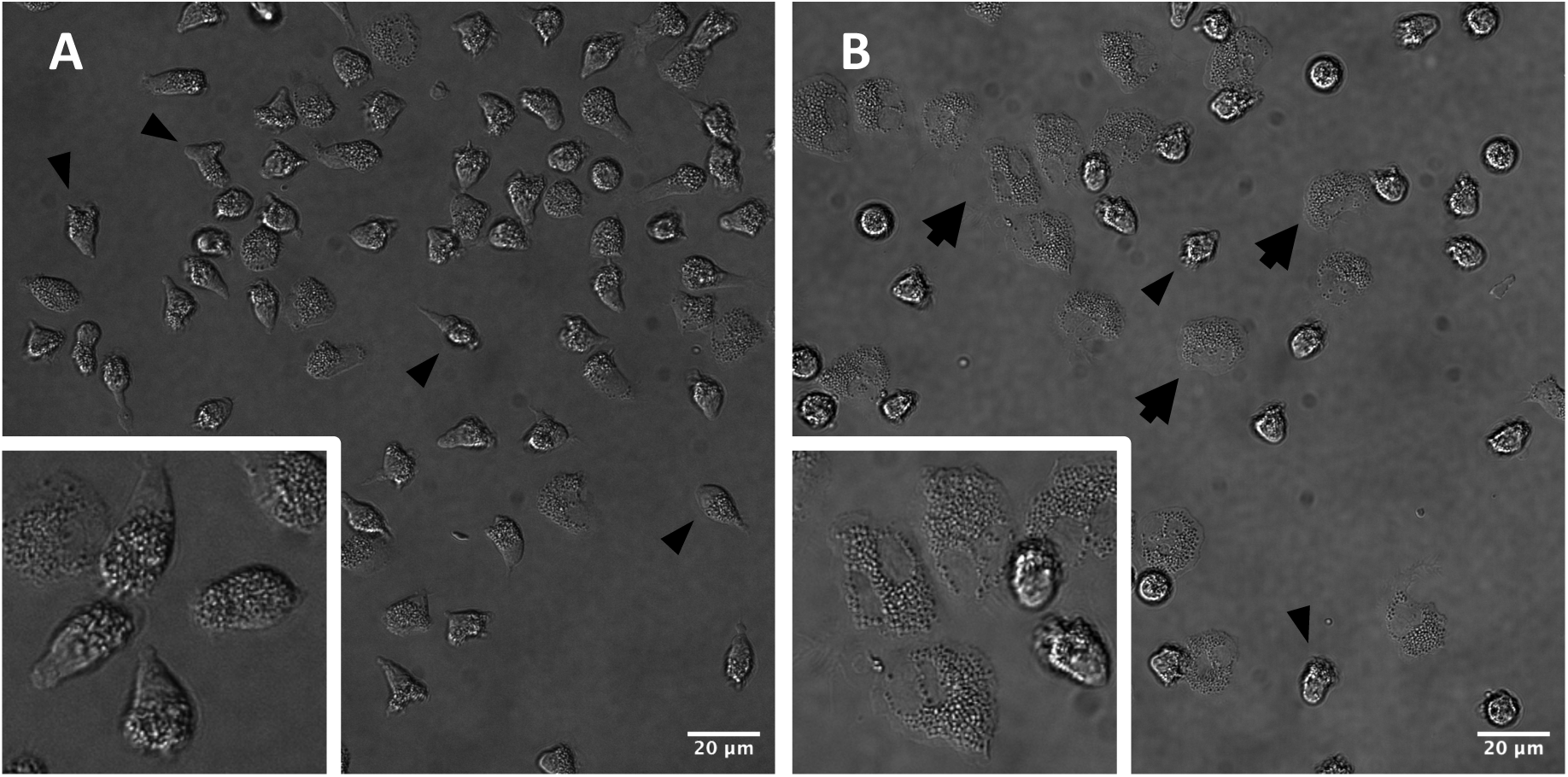
Morphologies of IL5- and IL33-activated eosinophils adherent to fibrinogen. (**A**) IL5-activated eosinophils adopt an acorn-like morphology (inset), with the nucleopod at the rear of the cell and the granulomere between the nucleus and leading edge. Arrowheads denote activated cells. (**B**) Morphologies of IL33-activated eosinophils evolve over time. Cells initially have a pear-like shape (arrowheads) into an intermediate that transitions into a flattened pancake-like cell (arrows, inset). Phase images captured on a Nikon A1R-Si+ confocal microscope. Single z-planes close to the substrate are shown.

Within 30 min, most (XX%) IL33-activated eosinophils flattened into a pancake-like shape with centralization of the nuclear lobes and dispersion of granules (**Figure 1**). Based on analysis of z-stacks, the height of IL33-activated eosinophils after 30 min was estimated to be 1-3 μm whereas the height of IL5-activated eosinophils was 5-7 μm (unpublished).

### Cytokine exposure and substrate define eosinophil cell fate

Loss of membrane integrity and cell death was monitored for up to 90 min as assessed by loss of calcein-AM from the cytoplasm concomitant with nuclear staining with ethidium homodimer. Over this period, cells activated with IL33 on fibrinogen had more cytolysis than cells activated with IL5. (**Figures 6 and 7)**. In the experiment shown, for the first 30 min, IL5-activated cells had little to no apparent cell death, followed by an increase to 3% at 60 min and 8% at 90 min. This is less than IL33, where at 30 min, 4% of cells were dead, with an increase to 13% by 60 min and 22% by 90 min. Although the percentages of dead cells at 60 min to 90 min varied from experiment-to-experiment, in 4 of 5 experiments with eosinophils on a substrate of fibrinogen, there was more death with IL33 than IL5. This suggests that IL33 acts as a cytolytic cytokine and induces cell death after extended exposure to eosinophils.

Substrate also plays a role in how eosinophils spread and how many undergo cell death. At 30 min, IL33-activated eosinophils show greater spreading than IL5-activated cells, with IL33-activated cells having flatter morphologies when plated on periostin (**Figure 8**) as compared to fibrinogen (not shown). We also observed more cell death, with significant increases in the percentage of dead cells at 90 min when comparing eosinophils on periostin to fibrinogen. At 90 min, death of IL5-activated cells 41%, and of IL33-activated cells 53%. Again, there was considerable variation in cell death at 90 min, but in 2 of 3 experiments there was more death on periostin than fibrinogen. These changes possibly reflect a greater cytolytic potential of periostin over fibrinogen.

### Nuclear collapse preludes cell death in IL33-activated eosinophils

While imaging IL33-activated eosinophils, we observed cells undergoing cell death (**Figure 9**). We noted cell death occurred in activated eosinophils and not in cells that were unactivated. Starting around ∼60 min, the bilobed nucleus loses its spread morphology and begins to condense (**Figure 9A**). We see this in the intensity of the calcein-AM staining as the diffuse coloration of the nuclear lobes becomes more intense. Once the lobes begin to condense, we note that they begin to draw towards one another until they form a condensed circular nucleus. At this point, ethidium homodimer-1 staining begins to infiltrate the cell, and the calcein-AM signal quickly dissipates. A truncated time course (**Figure 9B**) shows the condensed lobes of the nucleus beginning to approach each other (70-72.5 min), resulting in a single-lobed nucleus (75 min) that loses calcein-AM staining (77.5 min) and acquires red staining (>80 min). We note that between 77.5 min and 80 min, the nuclear membrane appears to have dissolved, which leads to more dramatic staining of nuclear content by ethidium homodimer-1. We find the same mechanism of nuclear collapse in other IL33-activated eosinophils but lack the video evidence in IL5-activated eosinophils from a lack of imaged cell death in this population. However, we do see the same ethidium homodimer-1 staining in IL5-and IL33-activated eosinophils and hypothesize that the same mechanism underlies cell death in these populations.

### Activation of eosinophils with either IL5 or IL33 causes vimentin to undergo extensive rearrangement

One of the striking differences between eosinophils stimulated with IL5 or IL33 for 10 min is in vimentin phosphorylation (**Table 1**). Upregulation of phosphosites at Ser7, Ser8, Ser9, Ser38, and Ser72 suggest activation of cycling dynamics to promote active reconstruction of vimentin intermediate filaments^26–28^. However, less is known about C-terminal phosphorylation such as Ser420 and Ser430, which show large increases in phosphorylation upon IL33, but not IL5, activation. Vimentin rearrangement underwent a dramatic change in methanol-fixed IL5- or IL33-activated eosinophils on fibrinogen (**Figure 2**). In unactivated cells in suspension, vimentin had a web-like morphology around the cell periphery (**Figure 2A**). In the adherent eosinophils, vimentin was in dense strands around the nucleus. Vimentin is widely used as a marker of epithelial to mesenchymal transition (EMT), which correlates with the dramatic morphological changes and polarization that eosinophils experience when becoming activated^29^. It is important for maintenance of nuclear shape and integrity in migrating mesenchymal cells^30^.

**Figure 2:**
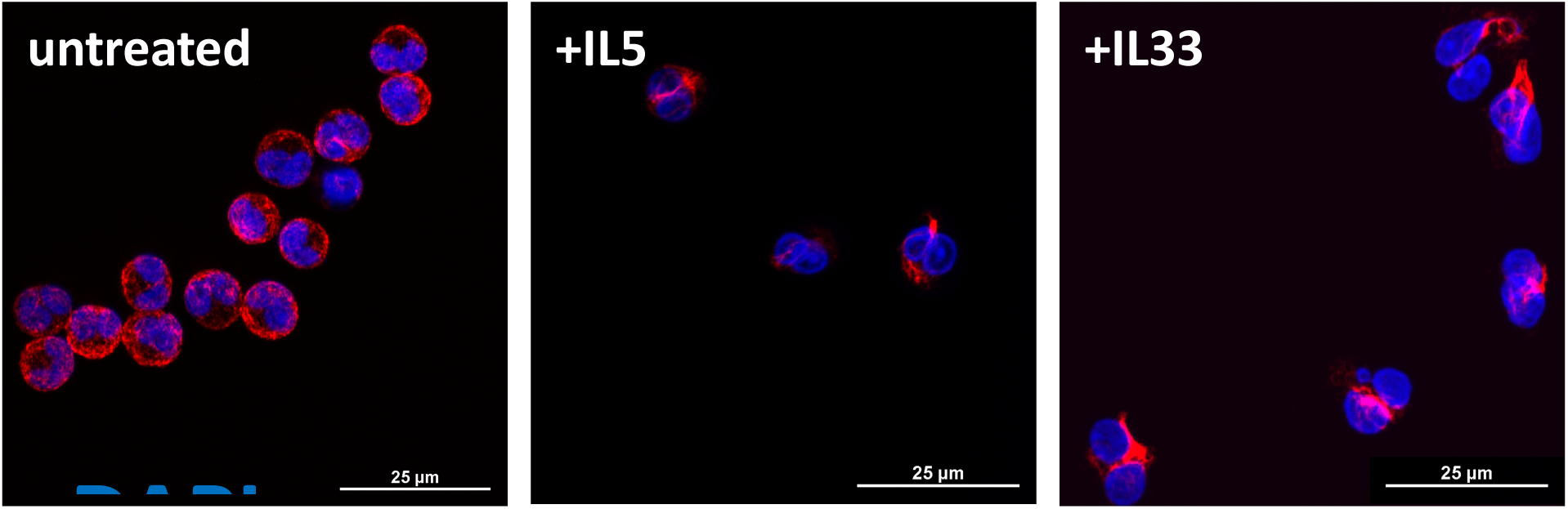
Vimentin condenses around the nucleus upon eosinophil activation. Unactivated eosinophils show a cage-like network of vimentin that surrounds the entire cell along the periphery of the membrane. IL5- or IL33-activated eosinophils adherent to fibrinogen show localization of vimentin to the nucleus, with greater intensity between nuclear lobes and at the tip of the nucleopod. Cells were fixed and permeabilized in ice-cold methanol. Fluorescence images captured on a Nikon A1R-Si+ confocal microscope. Maximum intensity projections are shown.

**Table 1:** Significant changes in phospho-sites of vimentin in activated eosinophils. Changes are color coordinated to cytokine activation: IL5 versus untreated, orange; IL33 versus untreated, green. For IL33 versus IL5, sites with more phosphorylation with IL5, orange; with IL33, green. Significance was defined as (fold change>|0.25|, q>0.05).

| Phosphosite | IL5 vs unactivated |  | IL33 vs unactivated |  | IL33 vs IL5 |  |
| --- | --- | --- | --- | --- | --- | --- |
|  | Present | Fold-change | Present | Fold-change | Present | Fold-change |
| Thr7 | X | -0.435 | X | 2.200 | X | -0.488 |
| Ser8 |  |  |  |  | X | -0.350 |
| Ser9 |  |  |  |  | X | 0.589 |
| Thr20 | X | -0.361 |  |  |  |  |
| Ser22 |  |  |  |  | X | 0.589 |
| Thr32 | X | -0.317 | X | -0.535 |  |  |
| Ser34 |  |  | X | -1.171 |  |  |
| Ser37 | X | -0.520 |  |  |  |  |
| Tyr39 | X | 0.481 |  |  |  |  |
| Ser42 | X | 0.771 | X | 1.126 |  |  |
| Ser47 | X | 1.239 | X | 0.530 |  |  |
| Ser49 |  |  | X | 1.204 |  |  |
| Ser50 |  |  |  |  |  |  |
| Ser55(Ser56) |  |  | X | 0.5121 |  |  |
| Ser71 |  |  |  |  |  |  |
| Ser72 | X | 1.071 | X | 1.171 |  |  |
| Ser73 | X | 0.825 | X | 0.419 | X | -0.365 |
| Ser412 |  |  | X | 1.425 |  |  |
| Ser420 |  |  | X | 3.113 |  |  |
| Ser430 | X | -0.800 | X | 2.444 | X | 1.683 |
| Ser436 |  |  |  |  | X | 0.494 |

**Table 2:** Donor characteristics and the performed experiments from purified eosinophils. A live and fixed imaging were done using eosinophils from 15 different donors. Substrate: fibrinogen (F), periostin (P). Experiment: Live/Dead imaging of activated eosinophils (Live/Dead), Imaging of SiR-tubulin and/or SiR-actin (SiR dyes), microtubule imaging with different permeabilizing agents (MP), vimentin imaging (VIM).

| Donor ID | Sex | Age | Asthma | Allergy | Activation | Substrate | Experiment | Live/Fixed |
| --- | --- | --- | --- | --- | --- | --- | --- | --- |
| BE0607 | F | 23 | Y | Y | IL33/IL5 | F/P | Live/Dead | Live |
| BE0628 | F | 53 | Y | Y | IL33/IL5 | F/P | Live/Dead | Live |
| BE0254 | M | 43 | Y | Y | IL33/IL5 | F/P | Live/Dead | Live |
| BE0660 | M | 31 | N | Y | IL33/IL5 | F/P | Live/Dead | Live |
| BE0670 | M | 37 | Y | Y | IL33/IL5 | F | Live/Dead | Live |
| BE0622 | M | 45 | Y | Y | IL33/IL5 | F | Live/Dead | Live |
| BE0527 | M | 33 | Y | Y | IL33/IL5 | F | Live/Dead | Live |
| BE0361 | M | 54 | Y | Y | IL33/IL5 | F | SiR dyes | Live |
| BE0622 | M | 45 | Y | Y | IL33/IL5 | F | SiR dyes | Live |
| BE0226 | F | 49 | Y | Y | IL33/IL5 | F | SiR dyes | Live |
| BE0659 | M | 51 | Y | Y | IL33/IL5 | F | SiR dyes | Live |
| BE0131 | M | 46 | Y | Y | IL33/IL5 | F | SiR dyes | Live |
| BE0165 | M | 50 | Y | Y | IL33/IL5 | F | MP | Fixed |
| BE0534 | M | 32 | Y | Y | IL33/IL5 | F | VIM | Fixed |

### Live imaging of microtubules reveals mesenchymal migration

Based upon the variable results imaging of actin and microtubules associated with different fixation and permeabilization conditions, we sought to image cytoskeletal structures without fixation. Silicon-Rhodamine (SiR) dyes that detect microtubules and f-actin are small molecules that bind between protomers of tubulin or actin proteins when polymerized. The associated dyes are not toxic and stable in living cells and can provide information about the dynamics of cytoskeletal structures as well as their localization in relation to other cellular features.

When SiR-tubulin dye was used to measure microtubule networks, previously unappreciated networks were visible. While a long-lived and stable population of microtubules emanating from the MTOC has been previously seen in static images^9,31^, the dye allows live-cell imaging. IL5-activated eosinophils had a bright population of microtubules that span the nucleopod from the tip to the MTOC tucked between the nuclear lobes. There also was a dynamic population of microtubules polymerizing and collapsing within the cytosol of the eosinophil, terminating within the granulomere and at the periphery of the cell at the leading edge; these underwent cycles of polymerization and collapse (**Figure 3**). The generation and collapse of these microtubules extending to the migratory front coordinates with cell movement, which suggests that eosinophil travel through mesenchymal migration^32,33^. This contrasts with another closely associated granulocyte, the neutrophil, which moves through amoeboid migration^34–36^.

**Figure 3:**
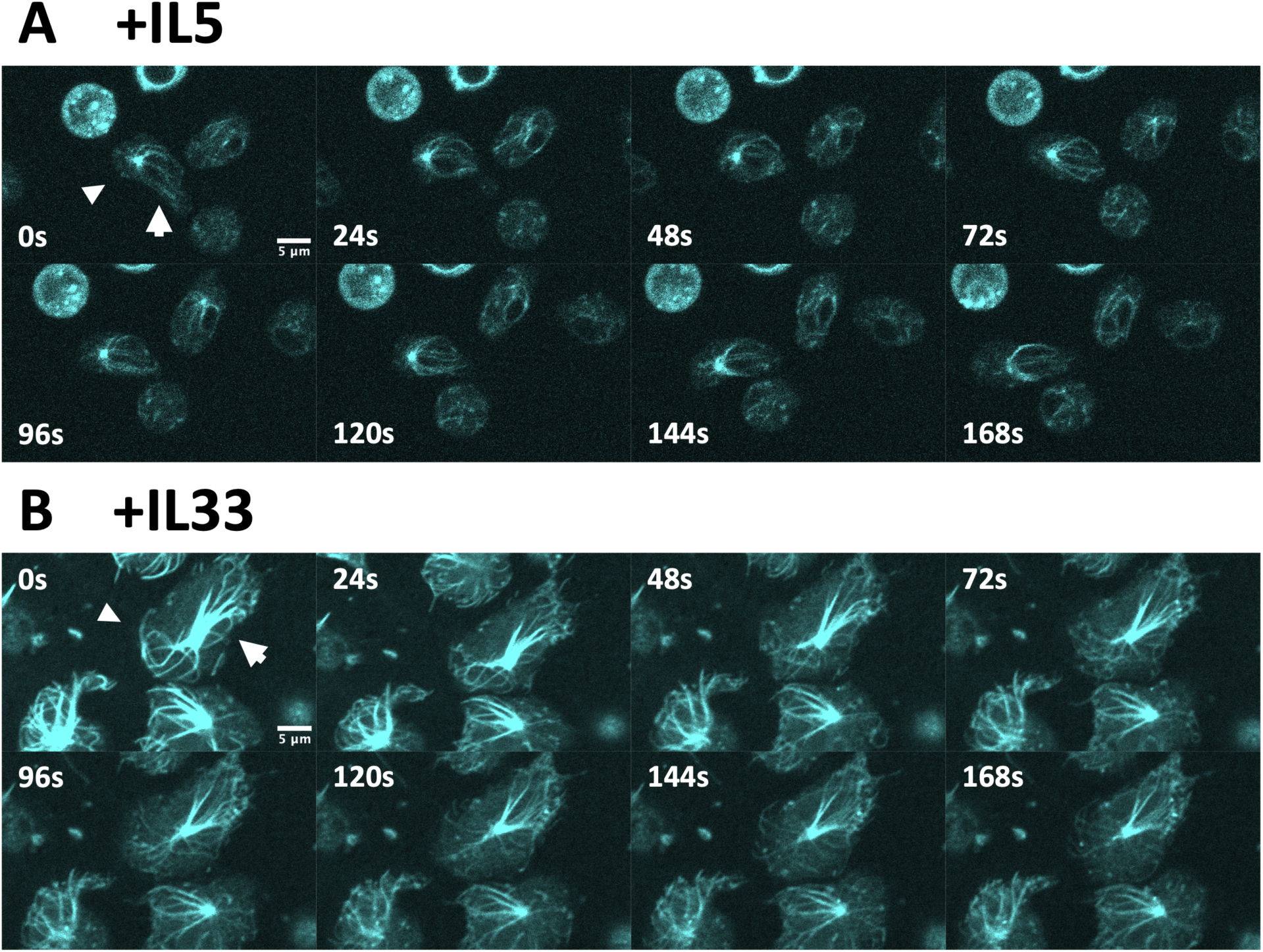
Live imaging with SiR-tubulin of microtubule networks in activated eosinophils on fibrinogen. (**A**) IL5-activated eosinophils are more migratory. The arrowhead marks the front of a cell where the more dynamic microtubule networks are located. The arrow marks the more stable microtubule network in the rear of the cell extending across the nucleopod. The bright dot found between these networks is the microtubule organizing center (MTOC). (**B**) IL33-activated eosinophils have a more dynamic network of microtubules at the cell periphery (arrowhead) and a dense stable network radiating over the nucleus from the MTOC. IL33-activated eosinophils are less motile, as seen across the set of images. Images captured on a Nikon A1R-Si+ confocal microscope and are single z-planes.

In migrating IL33-activated eosinophils, microtubules had the same localization as in IL5-activated cells, with a long-lived, bright population of microtubules that extend across the nucleopod, and a more dynamic and dimmer population that is found within the cytosol. The microtubule networks that coincide with cell movement sample the periphery of the eosinophil far more than IL5-activated cells, acting like an octopus sampling its environment with its tentacles. A bright spot found adjacent to the nucleopod and the granulomere corresponds with the centrioles found at the center of the MTOC (**Figure 4**). We see both a mother and daughter centriole, corresponding to the two bright spots seen in each centriole.

**Figure 4:**
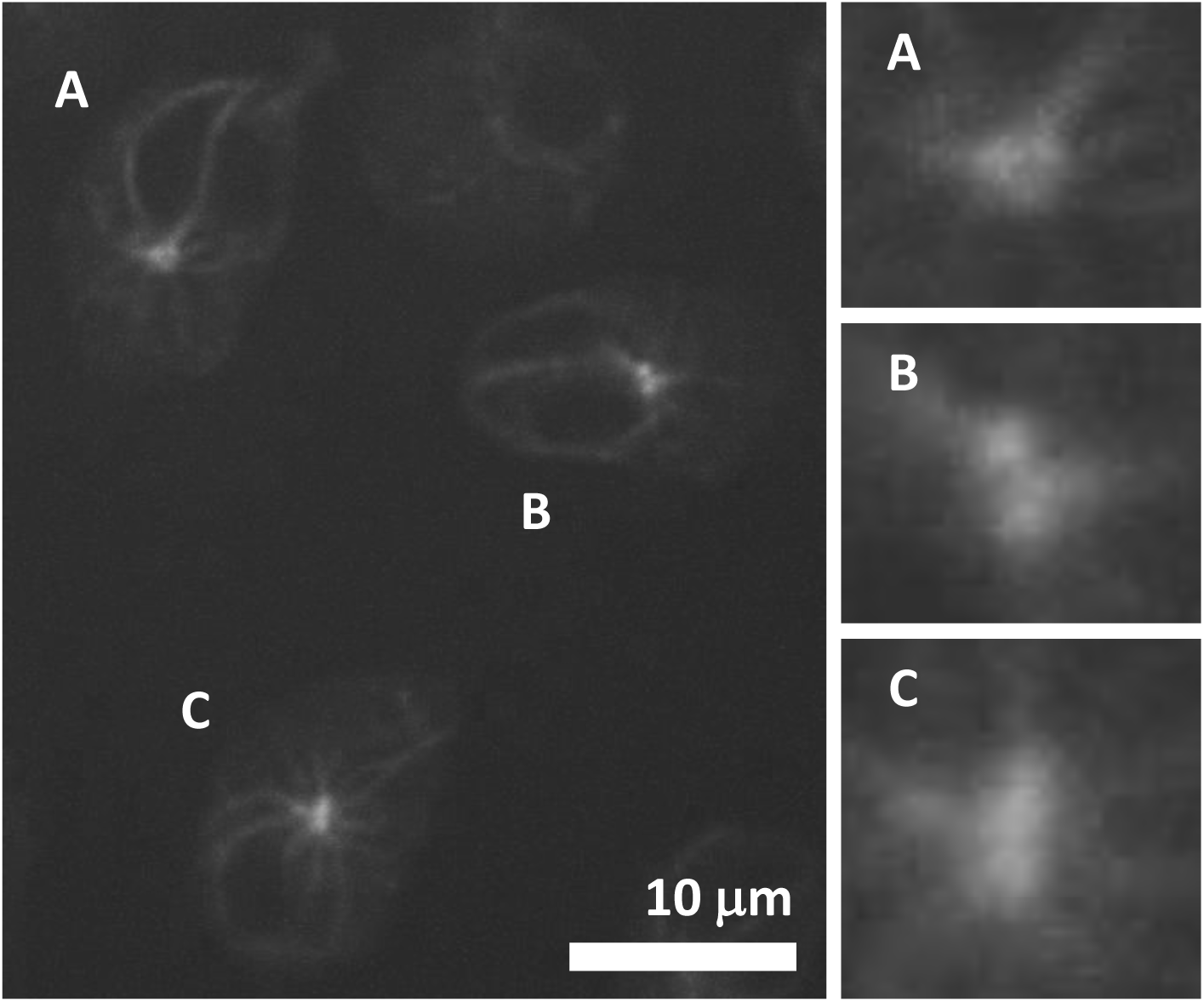
Mother and daughter centrioles at the MTOC of IL-5 activated eosinophils on fibrinogen. Two separate spots can be seen in three adjacent IL5-activated eosinophils (A, B, C) stained with SiR-tubulin. Microtubule networks can be seen radiating from the centrioles. Images captured on a Nikon W1 Spinning Disk confocal and are single z-planes.

### Live imaging of actin reveals f-actin at the nucleopod tip

SiR-actin showed a strong f-actin signal around the periphery of the cell and at the tip of the nucleopod (**Figure 5**). However, in contrast to previously reported imaging (Han paper), live imaging reports the strongest signal at the apical tip of the nucleopod. In IL5-activated eosinophils, the apical tip has focused f-actin staining within the entire tip, which seems to suggest an important structural feature that may be supporting this cellular protrusion. In IL33-activated eosinophils, the f-actin signal at the apical tip is more diffuse, as the tip itself is less defined. However, we still note a strong signal focused on a small region of the cell located at the end of the nucleopod. While neutrophils have a similar cellular feature, a uropod, at the rear of the cell, these cells show no similar staining, suggesting that this is a novel feature to cells with nucleopods^37–39^. A diffuse f-actin signal is seen within the granulomere, which suggests that the cytosolic content surrounding granules may include actin filaments.

**Figure 5:**
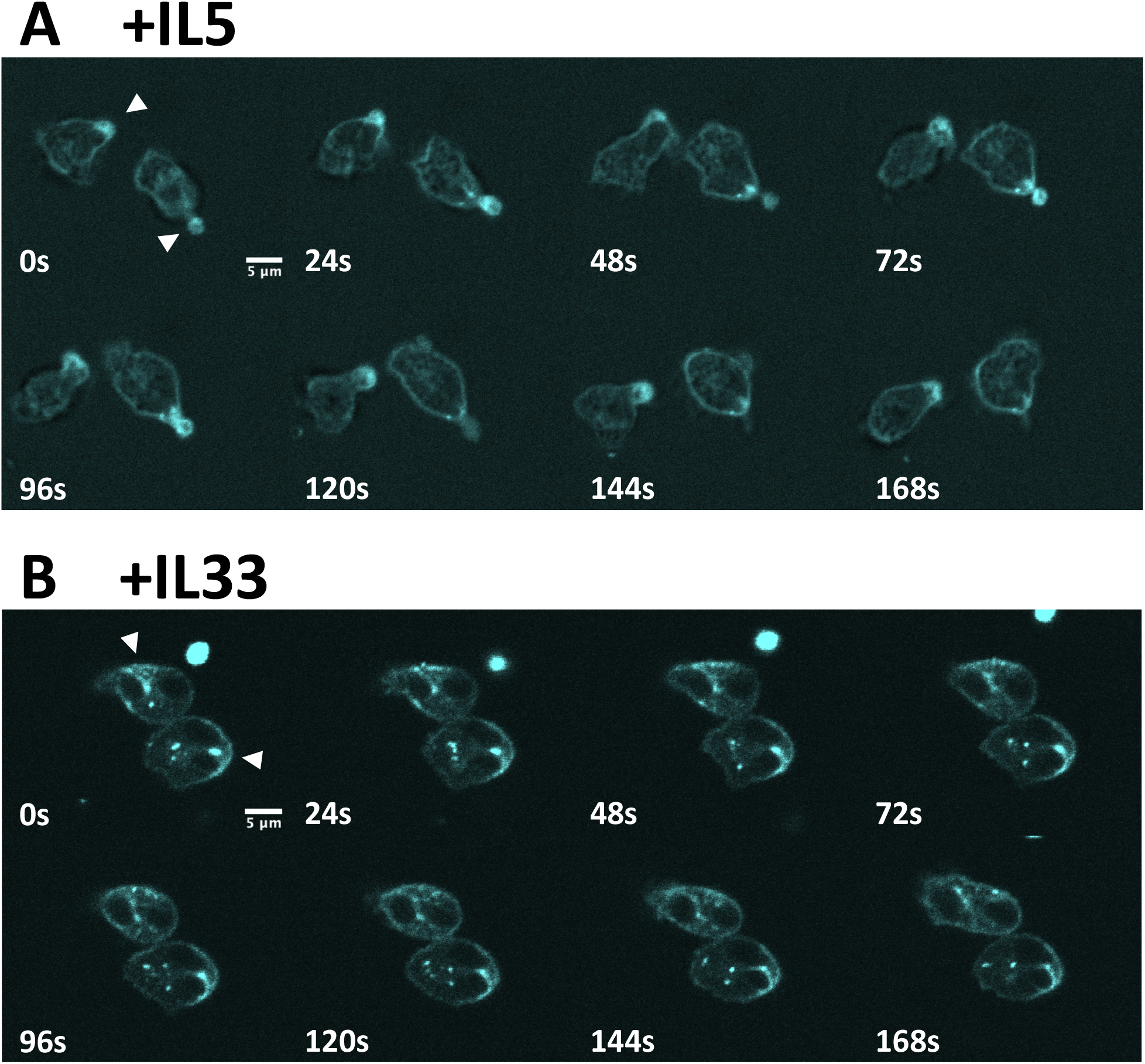
Live imaging with SiR-actin of f-actin networks in activated eosinophils on fibrinogen. **(A)** IL5-activated eosinophils are more migratory. Arrowheads denote the apical tip of the nucleopod, which shows bright f-actin staining. A ring of f-actin signal surrounds the cell, and a more diffuse f-actin staining is seen in the granulomere. **(B)** IL33-activated eosinophils have a less-defined apical tip with f-actin staining that is more intense than the rest of the periphery (arrowheads) as well as diffuse staining of the granulomere.

**Figure 6:**
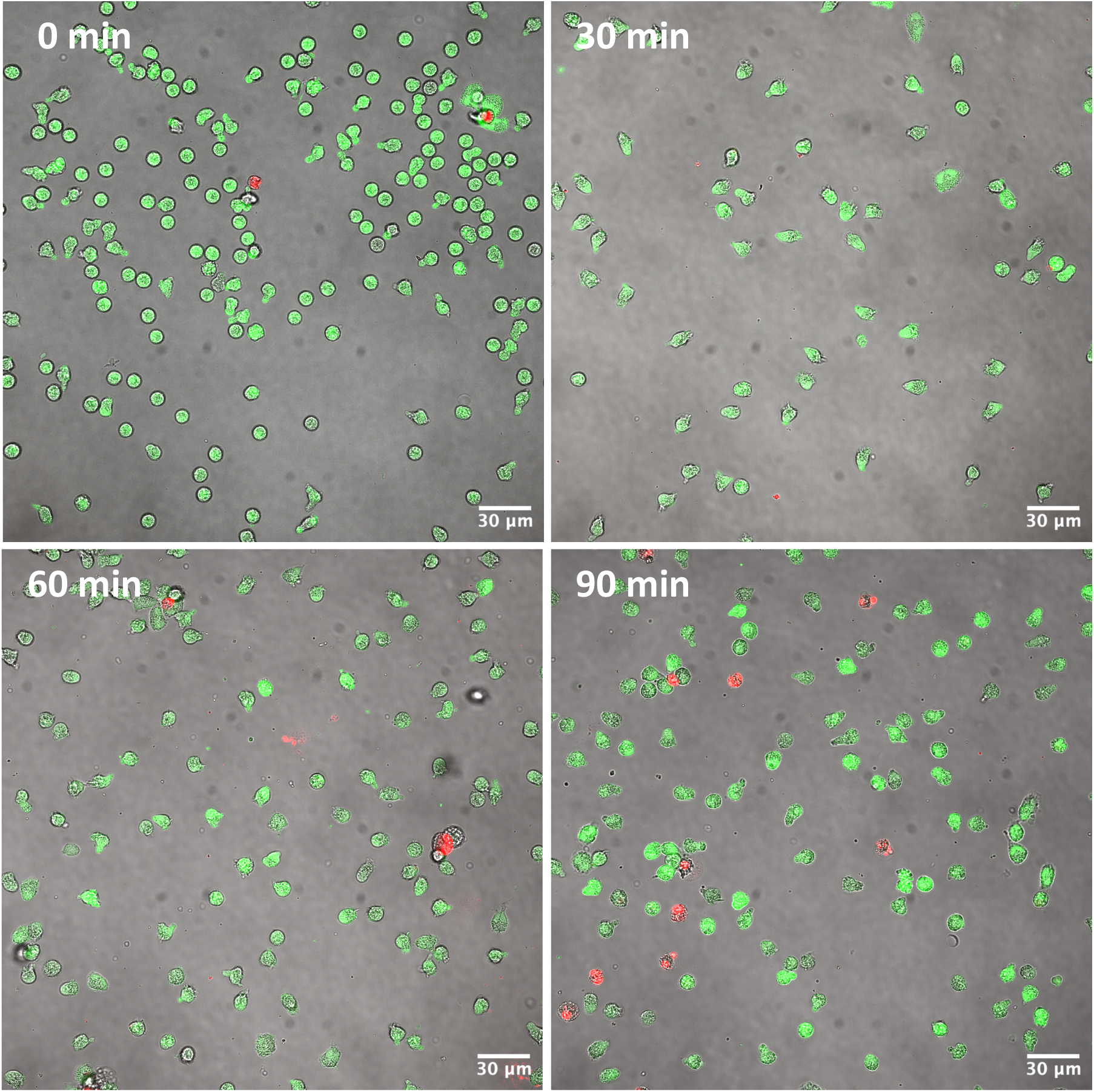
Live-dead imaging over 90 min of IL5-activated eosinophils on fibrinogen. Eosinophils were activated acutely with IL5 (50 ng/mL) and imaged in 30-min intervals to assess cell longevity. Cells progressively undergo cytolysis as assessed by loss of calcein-AM (green) from the cytoplasm and nuclear staining with ethidium homodimer (red). The percentages of dead cells are: 0 min, 0.1%; 30 min 0%; 60 min, 2.8%; and 90 min, 8.2%. Fluorescent images captured on a Nikon A1R-Si+ confocal microscope. Single z-planes are shown.

**Figure 7:**
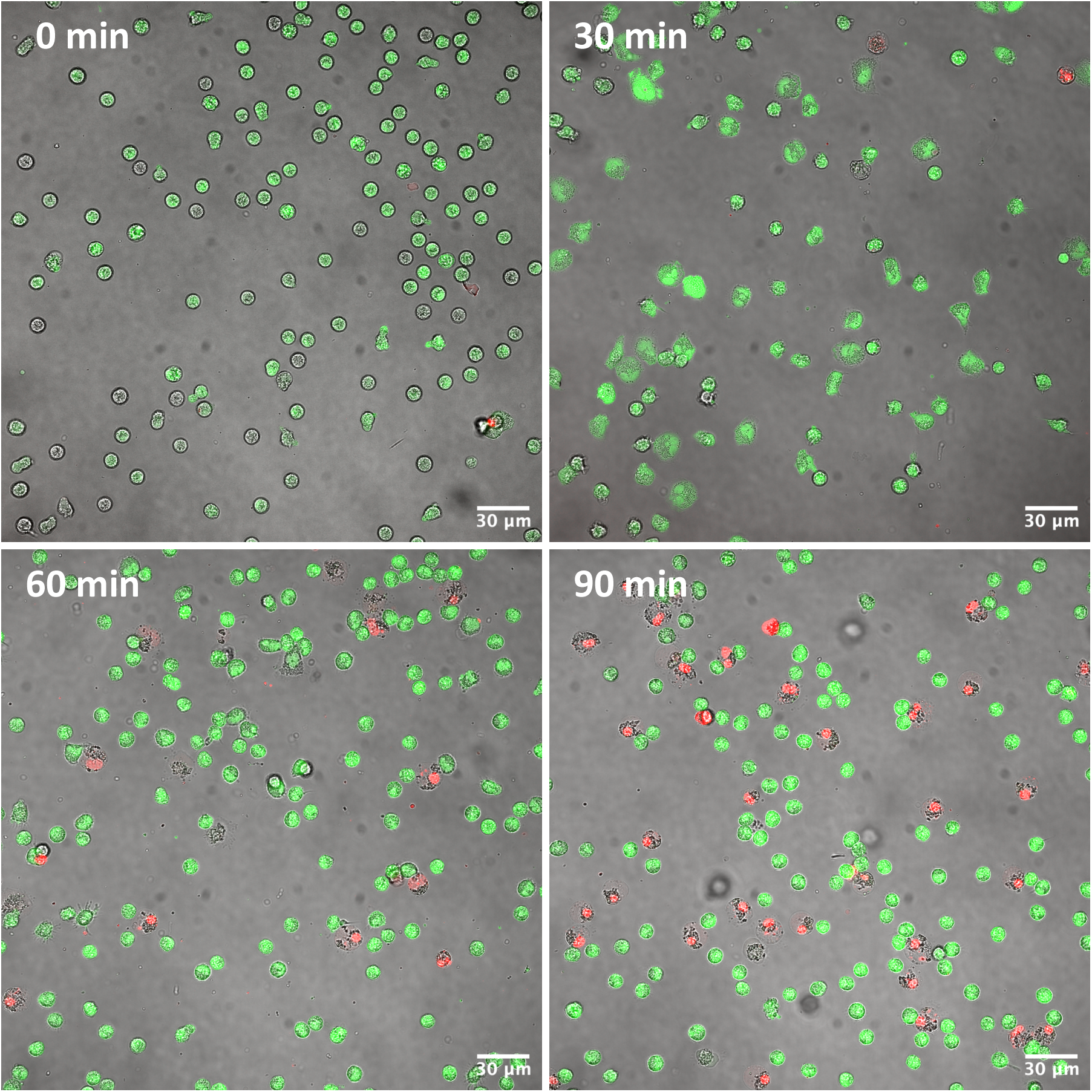
Live-dead imaging over 90 min of IL33-activated eosinophils on fibrinogen. Eosinophils were activated acutely with IL33 (50 ng/mL) and imaged in 30-min intervals to assess cell longevity. Cells progressively undergo cytolysis as assessed by loss of calcein-AM (green) from the cytoplasm and nuclear staining with ethidium homodimer (red). The percentages of dead cells are: 0 min, 0.7%; 30 min, 4.3%; 60 min, 12.8%; and 90 min, 22.2%. Fluorescent images captured on a Nikon A1R-Si+ confocal microscope. Single z-planes are shown.

**Figure 8:**
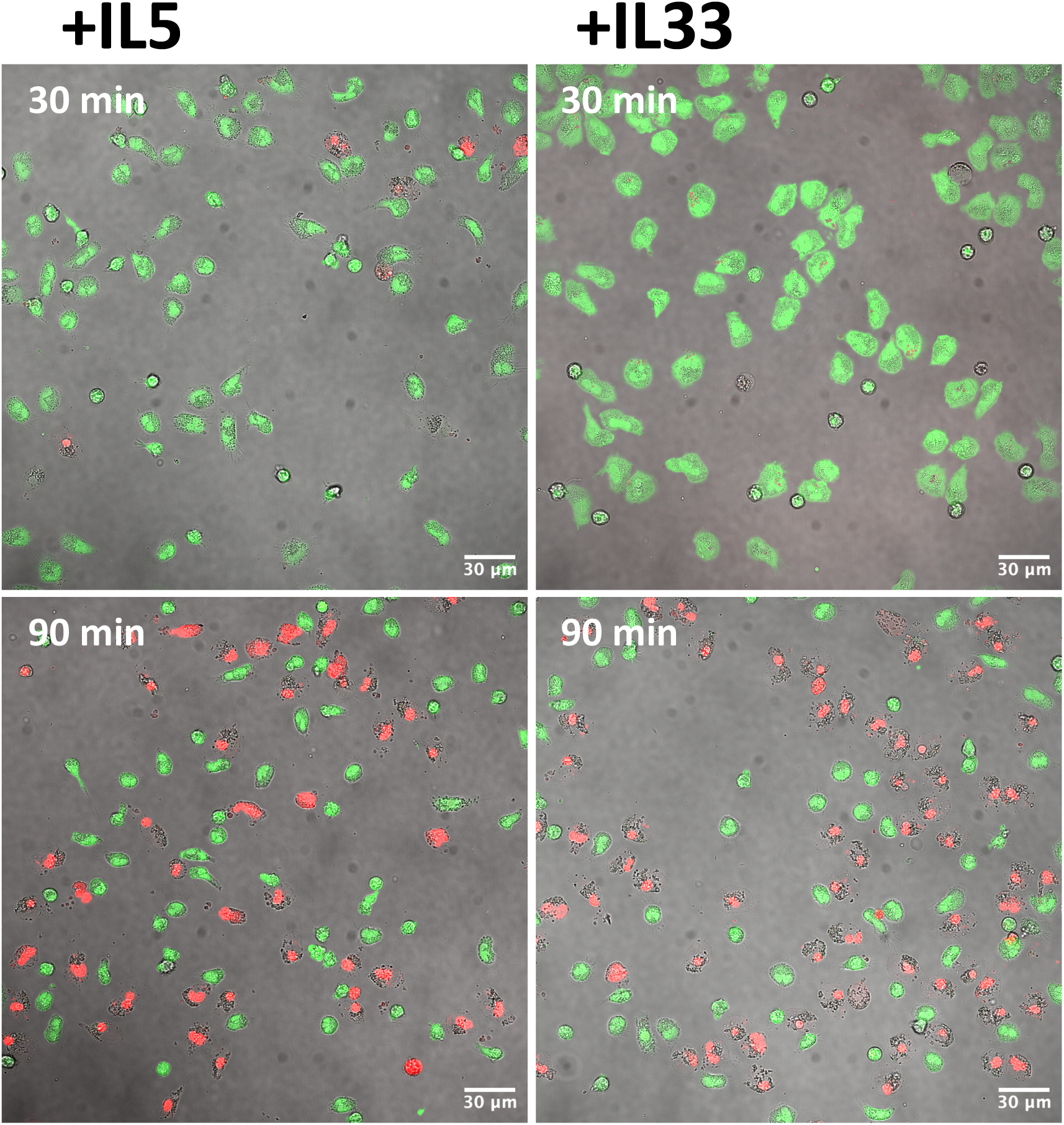
Live-dead imaging at 30 and 90 min of IL-5 or IL33-activated eosinophils on periostin. At 90 min, the percentages of cell death are: IL5, 40.9%; IL33-activated cells, 52.7%. Fluorescent images captured on a Nikon A1R-Si+ confocal microscope. Single z-planes are shown.

**Figure 9:**
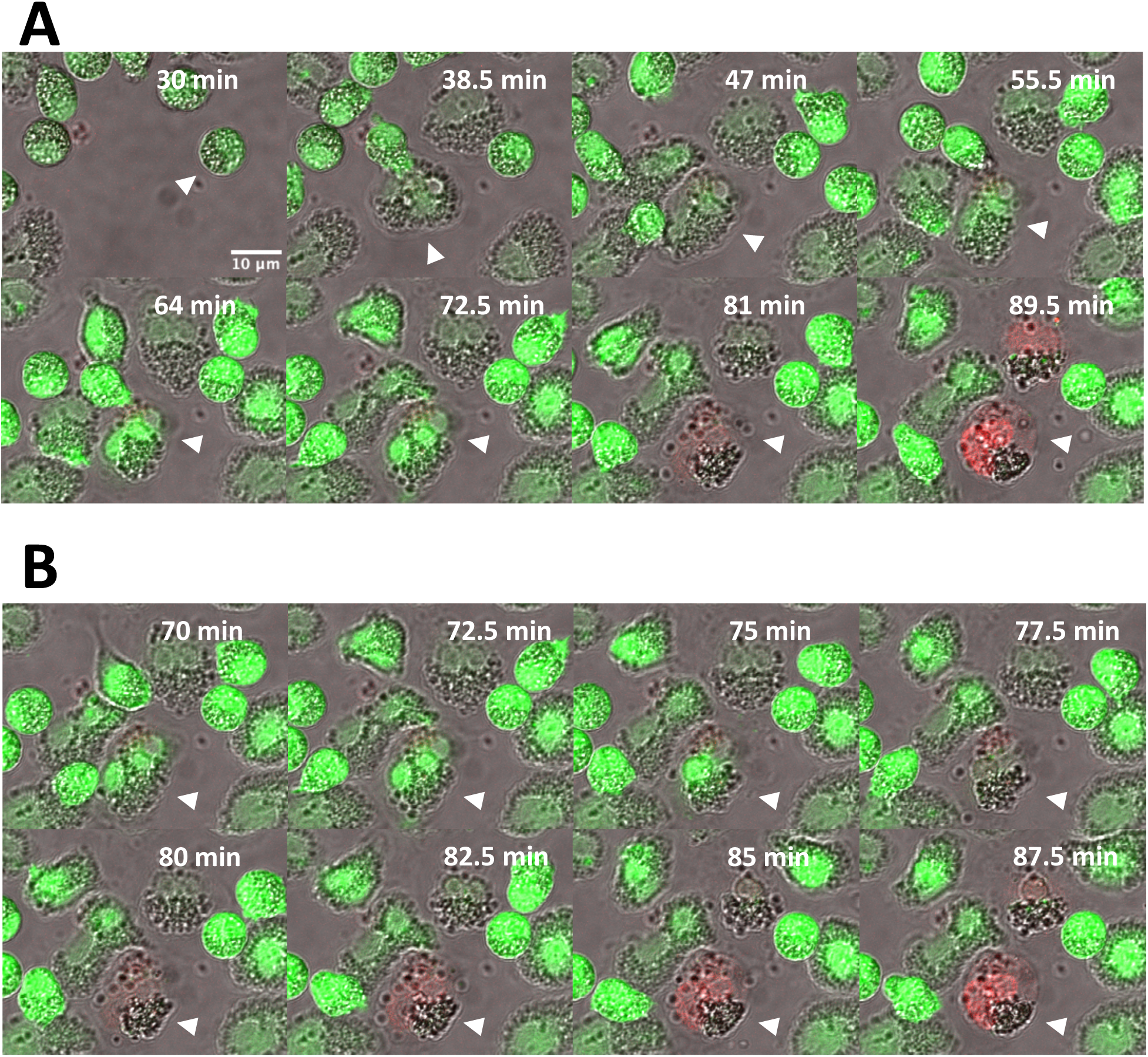
Live-dead imaging of IL33-activated eosinophils reveals nuclear integrity collapse and subsequent cell death. **(A)** An IL33-activated eosinophil (arrowhead) settles on to fibrinogen and spread. Starting around ∼60 min, the bilobed nucleus condenses into a single circular nucleus and dissipates, staining red. **(B)** A truncated time course shows the condensed lobes of the nucleus beginning to approach each other (70-72.5 min), resulting in a single-lobed nucleus (75 min) that loses calcein staining (77.5 min) and acquires red staining (>80 min). Fluorescent images captured on a Nikon A1R-Si+ confocal microscope. Single z-planes are shown.

## Discussion

*The response of eosinophils to IL33 has largely been uncharacterized. Previous work has shown that in solution, IL5 and IL33 show similar polarization of the eosinophil. For the first time, our live image demonstrates the significant morphological changes when activated eosinophil are exposed to an adherent substrate. Unlike IL5, which shows similar morphological polarization in solution and when adherent, IL33 activation is very different. In solution, IL33 appears more similar to IL5-activated eosinophils, having a pear-like morphology. This same morphology is present in IL33 activation on adherent substrate, but appears more as an intermediate state, as the eosinophils progress to a flattened pancake-like state upon prolonged adherence. However, how eosinophils may appear in complex environments, such as traversing*

Activation rearranges the cytoskeleton in eosinophils dramatically after activation within cytokine. Vimentin, which has previously been shown to be significant for nuclear stabilization in fibroblastic cells^30,40^, possibly has the same function in eosinophils, where the cage-like network condenses around the nucleopod once the cells become activated. This presumably is a prerequisite to extravasation, where nuclear integrity would become severally compromised without additional support.

Our research defines eosinophil migration as being mediated by microtubules, more closely resembling mesenchymal migration compared to amoeboid movement, which is mediated by actin. We see a robust network of microtubules extending to the migratory front of the cell, which coordinate with the motion of the eosinophil when observed by video microscopy. This contrasts with the amoeboid movement reported in neutrophils (Huttenlocher reference), in which f-actin staining is apparent at the migratory front. Live imaging of eosinophils has provided the first evidence that the extended network of microtubules that extends from the MTOC to the migratory front of the cell.

Together, this cytoskeletal imaging provides an updated image of the eosinophil (**Figure 10**). Upon activation, the cytoskeletal network shifts to prioritize a migratory morphology, with a vimentin cage that protects cell integrity focused to nucleopod and nuclear stability.

**Figure 10:**
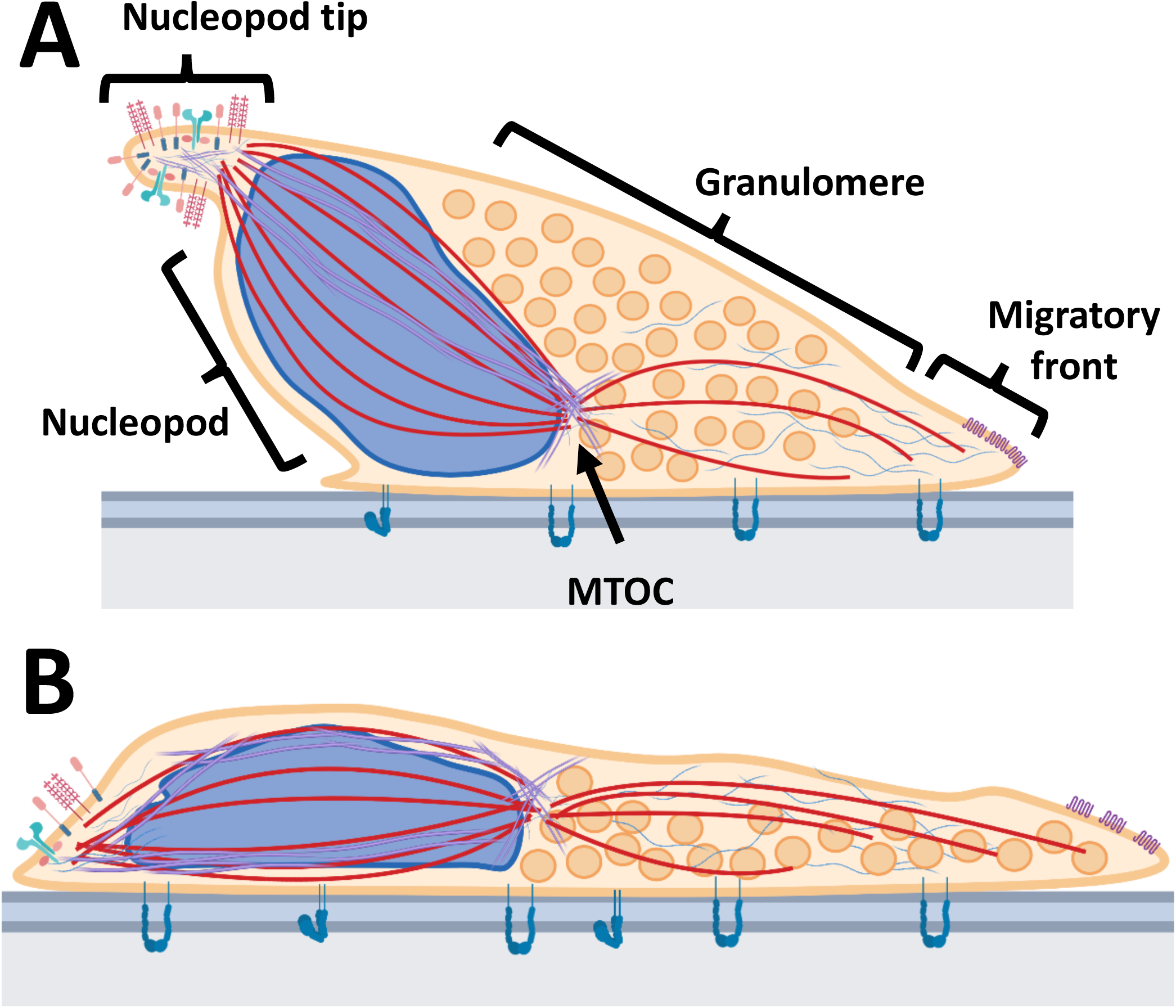
Cytoskeletal model of activated eosinophils. **(A)** The intracellular cytoskeletal model of an IL5-activated eosinophil and **(B)** an IL33-activated eosinophil. Our imaging highlights new features of the cytoskeleton of the activated eosinophil. Microtubules (red, thick line) traverse the nucleopod from tip to microtubule organizing center (MTOC) and from the MTOC to the migratory front of the cell. Vimentin (purple, double line) extends around the nucleus of the eosinophil. Actin microfilaments (thin blue squiggles) are found at the migratory front of the cell, granulomere, and tip of the nucleopod (labeled with various receptors).

Microtubule networks provide extensive networks that not only branch the nucleopod, but also direct migration of the cell with dynamic networks that rapid cycle to provide the mesenchymal migration we see in these cells^41,42^. This is coupled with actin, which is found at the cell periphery, staining the small lamellipodia that extend form the migratory front, drawing the cell forward^43^. The significance of actin within the tip of the nucleopod remains unknown and will require further study to elucidate its significance. In IL5-activated eosinophils (**Figure 10A**) we note greater migration of the eosinophils, with dynamic microtubule networks directing the movement of the cell. IL33-activated eosinophils (**Figure 10B**) are less motile and far thinner, with a single monolayer of granules creating a more diffuse granulomere. These observations corroborate previously published work in which IL33-activated eosinophils showed limited migration in respect to IL5-actiated eosinophils^24^. Microtubule networks also direct the movement of the cell, but the microtubules appear to be sampling the periphery of the cell more in this activated state. Overall, cytoskeletal elements of eosinophils experience dramatic reorganization, which in is similar in both IL5 and IL33 activated states. This suggests common features of reorganization that are generally required for eosinophils to become activated, leading to their subsequent adhesion and migration.

*We find that eosinophils strongly respond to stimulation with IL5 as a whole population, with >95% of cells becoming adherent, polarized, and becoming motile. However, the response to stimulation with IL33 has more nuance. When exposed acutely, IL33-activated eosinophils also respond with >95% of eosinophils taking on a polarized pear-like morphology, also showing the hallmark adherence, polarization, and motility seen in IL5-activated eosinophils. However, IL33-activated eosinophils can experience an additional transition to a flatter pancake-like morphology. Over the course of 30 mins, this population can become the predominant morphology of eosinophils, with 50-90% of cells belonging to this population.

Eosinophil survival based upon cytokine exposure and substrate was more thoroughly explored and defined in this work. Previous publications have detailed inflammatory nature of substrate assessing fibrinogen against murine and bovine collagen, but we sought to compare to human purified inflammatory substrates, fibrinogen and periostin^25^. Activation of eosinophils with IL5 maintains cell survival longer than IL33. Cells activated with IL33 had more spreading, less overall motility, and in the majority of experiments greater cell death at 90 minutes, suggesting that cellular response to IL33 may lead to cytolysis, whereas IL5 may act in part as a chemokine directing migration of cells. Substrate exposure also affects eosinophil longevity.

Eosinophils that were activated on fibrinogen-coated surfaces experienced fewer cell death events in both IL5- and IL33-activated eosinophils when compared to cells activated on periostin-coated surfaces, at equimolar coatings. We also note that IL5-activated eosinophils form a greater number of pancake-like cells on periostin than fibrinogen, which suggests that inflammatory substrates coordinate more adhesive responses in cells. This work highlights the significance of substrate exposure of eosinophils and the downstream cellular fate they commit to. While though fibrinogen has been noted to be a trigger of cytolytic degranulation in eosinophils, we demonstrate that periostin is a more potent trigger of this phenomenon^25^. *Further research is needed to dissect how ITAGAM receptor activation differs between these substrates, as well as how concentration dependency and substrate state define eosinophil response*.

Observed eosinophil death occurred more readily within IL33-activated eosinophils. We noted that pancake-like eosinophils more readily experienced cytolysis in comparison to pear-like IL33-activated cells, or acorn-like IL5-activate cells. Our observations align with previously imaged eosinophils, including observations made by electron microscopy *in vitro* and in tissue samples^44,45^. Vimentin has been noted as playing an essential role in maintaining the nuclear stability of migratory eosinophils (vimentin/eosinophil paper), and in IL33-activated eosinophils, we note drastic increases in phosphorylation at S420 and S430. The increased likelihood of cytolysis in this population may be a result of the phosphorylation of vimentin, the increased spreading of the nucleus and subsequent strain on the vimentin network to maintain nuclear stability, or a combination of these events. This observation suggests that coordination of nuclear integrity may play an important role in eosinophil fate and guide the timeline to cytolysis.

This work has significantly expanded the understanding of how cytoskeletal dynamics alter the morphological state of IL5- and IL33-activated eosinophils, and highlights the significant cytolytic role that IL33 plays when exposed to eosinophils^9^. No previous work has investigated the dynamics of cytoskeletal elements of fixed and live eosinophils on adhesive substrates. This work will help provide a more informed understanding of how cytokine exposure alters intracellular dynamics within eosinophils so that these cells can engage with substrate and become activated, and the function consequences of substrate recognition.

## Methods

### Purification of peripheral eosinophils from blood

Eosinophils were purified from 200 mL of heparinized blood from donors with allergic rhinitis or allergic asthma by Percoll centrifugation (density of 1.090 g/mL) and negative selection [e.g., ref. 21]. Donors had eosinophil counts in the high normal range of 200−500 per μL. The gradient separated eosinophilic and neutrophilic granulocytes from mononuclear cells such as NK cells and B cells. Negative selection with the AutoMACS system (Miltenyi, Auburn, CA) used a cocktail of anti-CD16, anti-CD14, anti-CD3, and anti-glycophorin A coupled magnetic beads.

Purity of eosinophils compared to other leukocytes was ≥98% as determined by Wright−Giemsa staining followed by microscopic scoring of cells and the viability was…≥x%. The studies were approved by the University of Wisconsin-Madison Health Sciences Institutional Review Board (protocol No. 2013-1570). Informed written consent was obtained from each subject before participation. Purified cells were kept on ice until being pelleted at 500x gravity for 10 min at room temperature, followed by resuspension in 37° C RPMI-1640 (+ L-glutamine, 25 mM HEPES, pH 7.4) supplemented with 0.1% human serum albumin (HSA). Eosinophils were rested for 1h in a tissue culture incubator under standard cell culture conditions (37° C, 5% CO_2_).

### Antibodies and other reagents

Eosinophils were stained with fluorescent dyes from the LIVE/DEAD Viability/Cytoxicity kit, for mammalian cells (Invitrogen, Waltham, MA) or SiR-Tubulin and -Actin dyes (Cytoskeleton, Inc., Denver, CO). Fixed cells were immunostained with mouse mAb anti-vimentin (V9) (Santa Cruz Biotechnolgy, Dallas, TX), nouse mAb antiα-tubulin (DM1A) (Sigma, St. Louis, MO), 4’,6-diamidino-2-phenylindole (DAPI, Life Technologies, Eugene, OR), rabbit control IgG (Danvers, MA), Alexa 647-conjugated phalloidin, Alexa Fluor 488-conjugated donkey anti-rabbit IgG, Alexa Fluor 555-conjugated donkey anti-mouse IgG secondary antibodies (Thermo Fisher Scientific, Madison, WI). Eosinophils were activated with recombinant human IL5 or IL33 (R&D Systems, Minneapolis, MN).

### Fixed imaging of adherent cells

Cells were unactivated or activated with IL-5 or IL-33 (50 ng/mL) at 37◦C. Following a 10 min incubation, cells were fixed and permeabilized via one of two methods: fixation and permeabilization with PFA and Triton X-100/SDS/digitonin/saponin/liquid nitrogen; or by fixation and permeabilization with the addition of ice-cold methanol for 5 minutes. After washing off methanol, coverslips were blocked in 10% BSA for 1 h and then incubated overnight at 4◦C in primary antibodies (Item II of Supplementary Data) diluted in 2% BSA and dilute detergent in PBS. The following day, coverslips were washed in PBS, and incubated for 1 h at room temperature with fluorophore-conjugated secondary antibodies specific for the species of the primary antibodies. After DAPI staining, coverslips were mounted on slides and sealed.

Images were acquired using a Nikon A1R-Si+ confocal microscope.

### Live imaging of adherent cells

Cells were applied to fibrinogen-coated (10 μg/mL) 35 mm MatTek dishes and incubated at 37◦C in a tissue culture incubator. Prior to activation eosinophils were incubated with: 150 nM SiR-actin or SiR-tubulin for 2h; 80 nM calcein-AM, 400 nM ethidium homodimer-1 Live/Dead dyes upon activation of eosinophils. Cells were activated with IL5 or IL33 (50 ng/mL) at 37◦C. Images were acquired using a Nikon A1R-Si+ confocal microscope with a motorized stage and Tokei Hit temperature-controlled chamber using a 40x/60x oil immersion objective at the University of Wisconsin-Madison Optical Imaging Core facility. 90 min videos with 1 frame/5s and individual images were acquired following activation. Videos and individual images were exported, using Nikon software. Videos were analyzed using Fiji software. Statistical analysis was done using GraphPad.

